# Aspen leaves as a “chemical landscape” for fungal endophyte diversity – Can nitrogen and herbivory shape the community composition in controlled conditions?

**DOI:** 10.1101/2020.12.26.424119

**Authors:** Johanna Witzell, Vicki Huizu Guo Decker, Marta Agostinelli, Carmen Romeralo Tapia, Michelle Cleary, Benedicte Riber Albrectsen

## Abstract

The endophytic microbiome may influence the ecological performance of plants, including forest trees. Various abiotic and biotic factors may shape the endophyte communities directly but also indirectly, by modifying the quality of host plants as a substrate. We hypothesized that potentially antifungal or fungistatic condensed tannins (CTs) would determine the quality of aspen (*Populus tremula*) leaves as a substrate for endophytic fungi. By subjecting the plants to nitrogen fertilization (N) or herbivory (H; leaf beetles) we aimed to change the internal “chemical landscape”, especially the CT levels, in aspen leaves. We expected that this would lead to changes in the fungal community composition, in line with the predictions of heterogeneity-diversity relationship hypothesis. To test this we conducted a greenhouse study where aspen plants were subjected to N and H treatments, individually or in combination. The chemical status of the leaves was analysed using GC/MS (114 metabolites), LC/MS (11 phenolics) and UV-spectrometry (CTs) and the endophytic communities were characterized using culture-dependent sequencing. Nitrogen treatment, alone or in combination with herbivory had a suppressing effect on the concentration and within-treatment variation in the CT precursors, catechins, and resulted in similar trend also in CTs. Nitrogen increased the concentration of certain amino acids, and it also tended to increase most of the analysed sugars sugars. Herbivory had a negligible effect on chemical traits. In N-treated plants, the endophyte richness and abundance were higher than in plants exposed to H, but in general, the diversity of the culturable endophytes remaind stable despite the subtle changes in leaf chemistry.

## Introduction

In natural ecosystems and cultivated environments, communities of fungal endophytes form a hidden stratum of biodiversity, imbedded inside plant tissues (Wilson & Lindow, 1994; Bailey et al., 2005; Albrectsen et al., 2010). Currently, studies on endophyte communities are in the frontline of plant ecology (e.g., Hardoim et al., 2015) and interest in these communities is also boosted by the prospects of utilizing the biosynthetic capacities of the endophytes in plant protection and plant growth promotion in agriculture and forestry (Busby et al., 2016; Witzell & Martín, 2018; Terhonen et al. 2019; Blacutt et al., 2020), as well as in medical or industrial applications (Nisa et al., 2015; Gouda et al., 2016). Advancement of molecular tools has made it possible to obtain more detailed information about the taxonomic structure of these hidden communities (Jumpponen & Jones 2009; Albrectsen & Witzell, 2012; Unterseher et al., 2016), but investigations are still challenged by temporal and spatial dynamics, the taxonomic complexity of the fungal community, and the lack of knowledge about the ecological roles of fungal species and their inter- and intraspecies interactions inside the plants (Witzell et al. 2014). Especially in the case of large-sized and long-lived forest trees the mechanisms determining endophyte community composition have remained puzzling (Ragazzi et al., 2003; Albrectsen & Witzell, 2012; Witzell & Martín, 2018).

The structure of endophyte communities has rarely been examined through theoretical frameworks of community ecology that aim to explain species diversity, abundance, and composition of species in communities, and the factors and processes underlying these patterns. Vellend (2010) states that all mechanisms that underpin patterns in ecological communities involve only four distinct processes, namely dispersal (movement of organisms across space); drift (stochastic changes in species abundance); speciation (creation of new species); and selection (which involves biotic and abiotic interactions including intra- and inter-specific interactions). The theoretical approaches to community ecology have been developed mainly for landscape level processes and for other organisms than fungi, but especially the selection process can be applied to the micro-environment inside plants. One of the central hypotheses in community ecology is that environmental heterogeneity promotes species richness, because it increases opportunities for niche partitioning (Ben-Hur & Kadmon, 2020). Translated to the spatial scale of a single plant, it predicts that a spatially or temporally heterogenic, internal chemical environment could support a more diverse endophyte community, and vice versa (a more uniform chemical environment would offer fewer nutritional niches).

In forest trees, endophyte communities are likely to be shaped by numerous environmental and host plant internal factors and processes. Trees receive their endophytes mainly from the surrounding environment (horizontal spreading; Clay & Holah, 1999; Petrini et al., 1993), and the influence of season and climatic conditions on the community structure has been found to be high (Gomes et al., 2018). Edaphoclimatic factors can strongly determine the dispersal and establishment of endophytes in trees (Carroll, 1995; Sieber, 2007), either directly by influencing the quality and quantity of the available inoculum, or indirectly, through effects on the host plant as a substrate for the fungi. Although landing of viable fungal spores on potential host plants may occur stochastically, the chemical environment on and inside the plant is likely to act as a selective factor, supporting the germination and growth of some fungi and suppressing others. For instance, several studies have found a connection between endophyte infections and phenolic plant metabolites in trees (Bailey et al., 2005; Albrectsen et al., 2010; Martín et al., 2013; Albrectsen et al., 2018). These ubiquitous plant metabolites are potentially antifungal or fungistatic (Witzell & Martín, 2008), but some fungi may also be able to use them as a carbon source (Blumenstein et al., 2015). Soil fertility is known to influence production of phenolics in plants, and especially the availability of soil nitrogen tends to be negatively correlated with phenolic concentrations in plant tissues (Bandau et al., 2015). Phenolic metabolism is known to readily increase in response to stress, and several phenolics have been suggested to have antioxidant, antimicrobial or anti-herbivore properties (Witzell & Martín, 2008; Gourlay & Constabel, 2019). Because phoretic associations between fungi and insects are likely to be common (Lewinsohn et al., 1994; Rice et al., 2007), insect herbivores may influence the fungal community in host tissues by acting as carriers of the fungal inoculum (Albrectsen et al., 2018). On the other hand, endophytic fungi may also influence the herbivores (Coblenz & Van Bael, 2013), further complicating the interpretation of interactions.

The goal of our study was to increase the current understanding of how foliar endophyte assemblages in woody plants vary in response to multiple, interacting factors. Our basic hypothesis was that the intrinsic status of phenolic metabolites, dominant defensive chemicals in most boreal and temperate zone trees species, would strongly influence the quality of plants as a substrate for endophytic fungi, and thus shape their community structure (Agostinelli et al., 2018; Albrectsen et al., 2018). Moreover, we anticipated that modification of the phenolic status using abiotic or biotic factors would result in changes in the fungal community. To test this hypothesis, we isolated endophytic fungi from leaves of vegetatively propagated plants of twelve aspen (*Populus tremula* L.) genotypes that were subjected to nitrogen amendment and leaf beetle herbivory, alone and in combination. The genotypes represented a similar chemotype in terms of their content of salicinoid phenolic glycosides (SPGs, low molecular weight phenolics that are characteristic for Salicaceae plants; Keefover-Ring et al. 2014), but differed with regards to concentration of high-molecular weight phenolics, condensed tannins (CTs; Robinson et al. 2012). The plant material thus allowed us to specifically focus on the importance of the latter compounds, which have earlier been identified as potentially antifungal compounds in aspen (Ullah et al. 2017).

We expected that fertilization would generally reduce the concentrations of phenolic metabolites and particularly CTs (Witzell & Shevtsova, 2004; Bandau et al., 2015; Decker et al., 2016) in aspen leaves. Furthermore, we anticipated that it would also reduce the within-group variation in phenolic concentrations, i.e. the phenolic concentration would vary less between N-treated replicate plants as compared to the control plant group (Edenius et al., 2012; Albrectsen et al., 2018). We expected that this quantitative response could lead to a more homogenous nutritional niche that would support a less diverse endophytic community than what is found in control plants. In contrast, we hypothesized that herbivory could lead to locally induced accumulation of constitutive or induced phenolics (Sampedro et al., 2011; Massad et al. 2014), or oxidation of phenolics due to tissue damage (Constabel et al. 2000; Salminen & Karonen, 2011). We expected that this would result in a more variable (multi-directional) and compartmentalized chemical environment in leaf endosphere, leading to higher diversity of nutritional niches and thus more diverse fungal communities, as predicted by the heterogeneity-diversity relationship hypothesis (Ben-Hur & Kadmon, 2020). The possible phoretic interactions could add to the diversity of fungal communities in H treated plants, especially in the strictly controlled greenhouse environment with limited environmental inoculum. We used the classical isolation method and molecular identification based on ITS1 and ITS2 regions that are widely used as the standard DNA barcode marker for fungi (Badotti et al. 2017) to describe the endophyte community. Despite the well-known limitations of this approach we expected it to be robust enough to capture the fast-growing fraction of the total fungal community that would most readily respond to the treatment effects in our short-term experiment. To study the metabolic status of the plants, we conducted a global metabolite analysis using GC/MS and completed it with a targeted LC/MS analysis of low-molecular weight phenolics. We also analyzed the concentrations of condensed tannins. The results are discussed within the framework of the heterogeneity-diversity relationship hypothesis (Ben-Hur & Kadmon 2020).

## Material and Methods

### Plant material

Plants belonging to twelve aspen (*Populus tremula* L.) genotypes were chosen from the SwAsp collection (genotypes number 4, 6, 7, 26, 41, 51, 69, 72, 79, 92, 98, and 100, Robinson et al. 2012). The selected clones represented the tremuloides-like chemotype, with four dominating SPGs (salicin, tremuloidin, salicortin and tremulacin) (Robinson et al. 2012; Keefover-Ring et al., 2014). Aspen plants were produced from in vitro tissue cultures (Umeå Plant Science Centre, Umeå, Sweden) and planted into 5 L pots in May 23th 2014. The pots where placed in the greenhouse 20°C, 60% R.H., and a 16:8 h D/N cycle. Lateral branches were removed within the first four weeks after planting to promote apical growth.

### Experimental set-up

Two individual plants from each of the 12 genotypes were randomly assigned to one of four treatments: control (C), nitrogen fertilization (N), herbivory (H), and their combination (NH) (n=24 plants per treatment). The whole experiment thus comprised a total of 96 plants. For N treatment, a Weibulls Rika-S^®^ solution (containing 84gr/liter nitrogen, NH_4_NO_3_) was applied weekly starting from June 17^th^ for four weeks. This resulted in a final N input that corresponds to the level of industrial forest fertilization in Sweden, 150 kg N ha^-1^ y^-1^ (Decker et al., 2016; Nohrstedt, 2001). The first fully expanded leaf on each plant was marked on June 17^th^ and this leaf was later used to assess baseline leaf chemistry (see below).

After four weeks of fertilization treatment, a mousseline fabric net was placed over the first three fully expanded leaves that had developed above the marked leaf after the first fertilization event. Five adult aspen leaf beetles (*Chrysomela tremula* Fabricius) were placed into the nets after the last fertilization event and allowed to feed for five days (treatments H and NH). The beetles had been reared in the laboratory at Umeå Plant Science Centre since the beginning of May and belonged to the second generation of a culture captured close to Ekebo, in southern Sweden. Empty nets were placed in similar position also on plants in C and N treatments to control for the effect of the mousseline bag.

### Sampling

After five days of herbivory treatment, the experiment was ended. Leaves were collected from each plant for phytochemical analysis (the leaf below the netted leaves) and for fungal endophyte analysis (the netted leaves). All the leaves were morphologically at the same stage, i.e., fully expanded, and thus expected to have comparable activity of secondary metabolism. Leaves for the chemical analyses were flash-frozen in liquid nitrogen, freeze-dried and stored at −20°C until they were ground to fine powder using a bead mill (MM 301 Vibration Mill, Retsch GmbH and Co. KG, Haan, Germany) at 25 Hz for 3 minutes. The fine powder was stored in vials at −20°C until chemical analyses. Leaves designated for endophyte analysis were collected and stored at 4°C for a maximum of three days prior to surface sterilization and endophyte isolation. In case the petiole of a netted leaf was chewed by the beetle (causing wilting or the death of the leaf), those leaves were discarded from further analyses. A total of 90 plants were included in the endophyte analysis. The growth of plants and insects was recorded (Table S1).

### Phytochemical analysis

For the global GC/MS analysis, 6.00 (± 1.00) mg of leaf powder was extracted in 1 ml methanol:chloroform:water (v:v:v) at 4 °C. Deuterated salicylic acid [^2^H_6_] (Isotec, Miamisburg, USA) was used as an internal standard in all samples. Samples were centrifuged at 4 °C and 100 μl of the resulting supernatant was evaporated in vacuum. The residues were resolved in 30 μl of methoxyamine (15 μg/μl in pyridine) and 30 μl of MSTFA. Methyl stearate (30 μl of 15 ng/μl in heptane) was added before analysis, and 1 μl of each aliquot injected by a CTC Combi Pal autosampler (CTC Analytics AG, Switzerland) into an Agilent 6890 gas chromatograph. The chromatograph was equipped with a 10 m x 0.18 mm fused silica capillary column with a chemically bonded 0.18 μm DB 5-MS UI stationary phase (J&W Scientific). The compound detection was performed in a Pegasus III time-of-flight mass spectrometer, GC/TOFMS (Leco Corp., St Joseph, MI, USA). MATLAB™ R2011b (Mathworks, Natick, MA, USA) was used to quantify the mass by means of integrated peak areas. All pre-treatment data procedures, such as base-line correction, chromatogram alignment, data compression and Hierarchical Multivariate Curve Resolution (H-MCR), were performed using custom scripts according to Jonsson et al., (2005). The extracted mass spectra were identified by comparisons of their retention index and mass spectra with libraries of retention time indices and mass spectra (Schauer et al., 2005). Identification of compounds was based on comparison with mass spectra libraries (in-house database) as well as Kovats retention index. The identified chemicals were quantified by peak area and assigned to metabolite classes (phenolics, amino acids, fatty acids) (Table S4).

For the targeted LC/MS analysis, samples were prepared as above, but the residue was suspended in 25 μl of methanol and 25 μl of 0.1% v/v aqueous formic acid. A 2.0 μl aliquot was injected onto a C18 UPLC™ column (2.1 × 100 mm, 1.7 μm) and chromatographic separation was performed on a LCT Pemier TOF/MS in negative mode (Waters, Milford, MA, USA), following the method by Abreu et al., (2011). The standard compounds used in this analysis were from our in-house library (the Swedish Metabolomics Centre: SMC, Umeå, Sweden) including salicinoid standards (salicin, tremulacin, salicortin, and tremuloidin). MassHunter™ Qualitative Analysis software package (version B06.00, Agilent Technologies Inc., Santa Clara, CA, USA) was used to acquire the Mass Feature Extraction (MFE) and extracted features were aligned and matched between samples using Mass Profiler Professional™ 12.5 (Agilent Technologies Inc., Santa Clara, CA, USA). The concentrations of salicinoids were quantified according to the peak area of each compound using linear standard curves. Masses of either or both of the deprotonated ion ([M−H]−) and the formate adduct ([M−H + FA]−) were assessed, based on molecular weights according to Abreu et al., (2011) and Keefover-Ring et al., (2014) and guided by retention times where available (Table S4).

Condensed tannin concentrations were assessed based on a modified Porter’s assay (acid: butanol method (Porter et al., 1985; Bandau et al., 2015): 10± 1 mg fine powder of freeze-dried leaf sample was added to 800 μl of an acetone/ ascorbic acid solution (70% acetone, 30% Milli-Q water, with 10mM ascorbic acid) then incubated with an iron and acid-butanol reagent in boiling water for 1h. Absorbance was measured on a Spectra Max 190 microplate reader (Molecular Devices, Sunnyvale, CA) at A 550 nm. Procyanidin B2 (C_30_H_26_O_12_, Sigma-Aldrich^®^, St. Louis, MO, USA) was used as the tannin standard (Table S4).

### Fungal endophyte isolation and identification

Endophytic fungi were isolated following the method described by Albrectsen et al. (2010). From each leaf (three leaves per plant in C and N treatments; one to three leaves per plant in H and NH treatments, a total of 228 leaves), ten segments (about 1×1 cm) were cut using a flame-sterilized scalpel under aseptic conditions (in the laminar flow bench). The segments were surface-sterilized using 95% ethanol and 2% sodium hypochlorite, and rinsed in sterile water before placing them individually on potato dextrose agar (PDA-agar; Sigma-Aldrich^®^, St. Louis, MO, USA) amended with streptomycin (12×12 cm dishes), as described in Albrectsen et al. (2010). A thin strip of Parafilm was used to keep the lid of the dishes on place. The cultures were checked for fungal growth every second day for one month. Emerging colonies were transferred to new Petri dishes (PDA) to obtain pure cultures. The plates were opened only inside the laminar, the fungi were transferred using flame-sterilized tools, and utmost care was taken to not contaminate the dishes with external fungi.

The recovered fungal cultures (n=1642) were classified into 30 distinct morphotypes (MTs) based on their visual morphological traits and growth rate estimated on a four step scale (fast, medium, slow, very slow). In order to obtain information about the taxonomy of the fungal community, a total of 59 cultures representing the 30 MTs were selected for sequencing. From each culture, plugs of PDA containing growing mycelium were transferred to malt extract broth. After an incubation period of two weeks (room temperature, darkness), mycelia were filtered, placed in 50 mL Falcon tubes (Fisher Scientific, Waltham, Massachusetts, US) and lyophilized for 48 hours. Freeze-dried samples were then pulverized in a FastPrep^®^-24 homogenizer (MP Biomedicals, Santa Ana, USA). Total genomic DNA was extracted using the E.Z.N.A. SP Plant DNA Kit (Omega Bio-Tek, Inc., Norcross, USA). The extracted nuclear DNA was measured using NanoDrop^®^ ND-1000 (Wilmington, USA). PCR amplification of the internal transcribed spacer (ITS) region of the rDNA was performed with the primers ITS1 and ITS4 (White et al., 1990). Each 50 μl PCR reaction mixture contained 5 μl of 10x PCR buffer, 0.4 μM of each primer, 0.2 mM dNTP, 1.5 mM MgCl_2_, 1U Taq polymerase and 10 ng fungal DNA. PCR program consisted of 94°C for 5 minutes; 33 cycles of 94°C for 30sec, 55°C for 30sec, 72°C for 30 sec and 72°C for 10 min. PCR samples were kept at 4°C before analyzing via gel electrophoresis on 1.5% Agarose (BIO-RAD) gel and visualized under UV light with 0.1% GelRed (GelRed™ Nucleic Acid Gel Stain, 10,000X in DMSO). PCR products were purified with the HT ExoSAP-IT High-throughput PCR Product Cleanup (Affymetrix, Santa Clara, USA) following manufacturer’s instruction. After quantification of DNA concentrations with Qubit fluorometer 3.0 (Life Technologies, Carlsbad, USA), samples were sent to the National Genomics Infrastructure (NGI) at Science for Life Laboratory (SciLifeLab, Uppsala, Sweden) Visualization of raw sequence data was done with the software Chromas (version 2.4.4, Technelysium, South Brisbane, Australia). All sequences were aligned and manually edited using BioEdit (Ibis Biosciences, Carlsbad, CA, USA). Sequencing data were blasted against best matches in the reference database at NCBI (http://www.ncbi.nlm.nih.gov/). The ITS sequence homology for delimiting fungal taxa was set to >98.5% for presumed species (Table S2).

### Data analysis

The Kruskall-Wallis test was used to test the treatment effects on specific chemicals and statistical analyses. The images including PCA plots (for 114 targeted and non-targeted metabolites from LC/MS and GC/MS) and 11 targeted phenolics (mainly SPGs from LC/MS) were generated with help of R version 3.6.1 (2019-07-05) packages: ggplot2 (https://cran.r-project.org/web/packages/ggplot2/), FactorMineR (https://cran.r-project.org/web/packages/FactoMineR/index.html) (Lê et al., 2008), and factoextra (https://cran.r-project.org/web/packages/factoextra/index.html).

Fungal diversity, (richness, S; Whittaker, 1972) was tracked as the number of distinct morphotypes (MTs) found in each plant, and the MT abundance (frequency) was recorded as the number of isolates of a given MT per sampled plant (n=24 plants for C and N, and 21 plants for H and NH treatments). Venn diagram was constructed to illustrate the number of shared and unique MTs per treatment (Shade & Handelsman; 2012). Relative abundance was determined for each MTs as the number of isolates of the given morphotype divided by the total number of isolates emerged from all samples (n=1642). To examine the possible shifts in the community composition among the different treatments (C, N, H and NH) a permutational multivariate analysis of variance (Permanova) of relative abundances was performed with the adonis function in vegan package (https://cran.r-project.org/web/packages/vegan/index.html) using 999 permutations, followed by a pairwise comparison with pairwise.adonis (Martinez, 2019).

MT richness values were used to create sample-size-based rarefaction (interpolation) and extrapolation (prediction) curves (Colwell et al., 2012, Chao et al., 2014, Hsieh et al., 2016) with an end point of 42 individuals and 100 bootstrap repetitions. The curves were generated with iNEXT (iNterpolation and EXTrapolation) R package (https://cran.r-project.org/web/packages/iNEXT/index.html) and visualized with ggiNEXT, the ggplot2 extension for iNEXT. To compare the complexity of communities, Simpson’s (*D*) and Shannon (*H*’) diversity indices and Pielou’s index for evenness (*J*’) were calculated using the Vegan package (Oksanen et al., 2016; Table S3). The effect of the treatments (C, N, H, NH) on the ecological diversity indexes and on the fungal richness (S) was assessed with an analysis of variance followed by a multiple comparison test (residuals were checked for normality, homoscedasticity and linearity). When the data did not meet these requirements, a robust ANOVA based on trimmed means was performed with WRS2 package (https://cran.r-project.org/web/packages/WRS2/index.html). Before all the analyses were performed, data was balanced using a decision tree to generate new samples from the minority classes (H and NH treatments) with an oversampling algorithm (random walk oversampling) from the imbalance package in R (https://cran.r-project.org/web/packages/imbalance/index.html).

## supplemental material

Supplementary file 1 (Tables S1, S2, S3; PDF), Supplementary file 2 (Table S4, Excel file).

## RESULTS

### Treatment effects on aspen chemistry

The score plot (PCA) of global metabolite analysis (data included 114 metabolites) showed a slight separation between fertilized and non-fertilized treatments (Fig 1a). The plot of targeted metabolites (11 phenolics, including catechin but excluding condensed tannins) did not show any clear distinction between fertilized and non-fertilized plants (Fig. 1b).

**Fig. 1.**
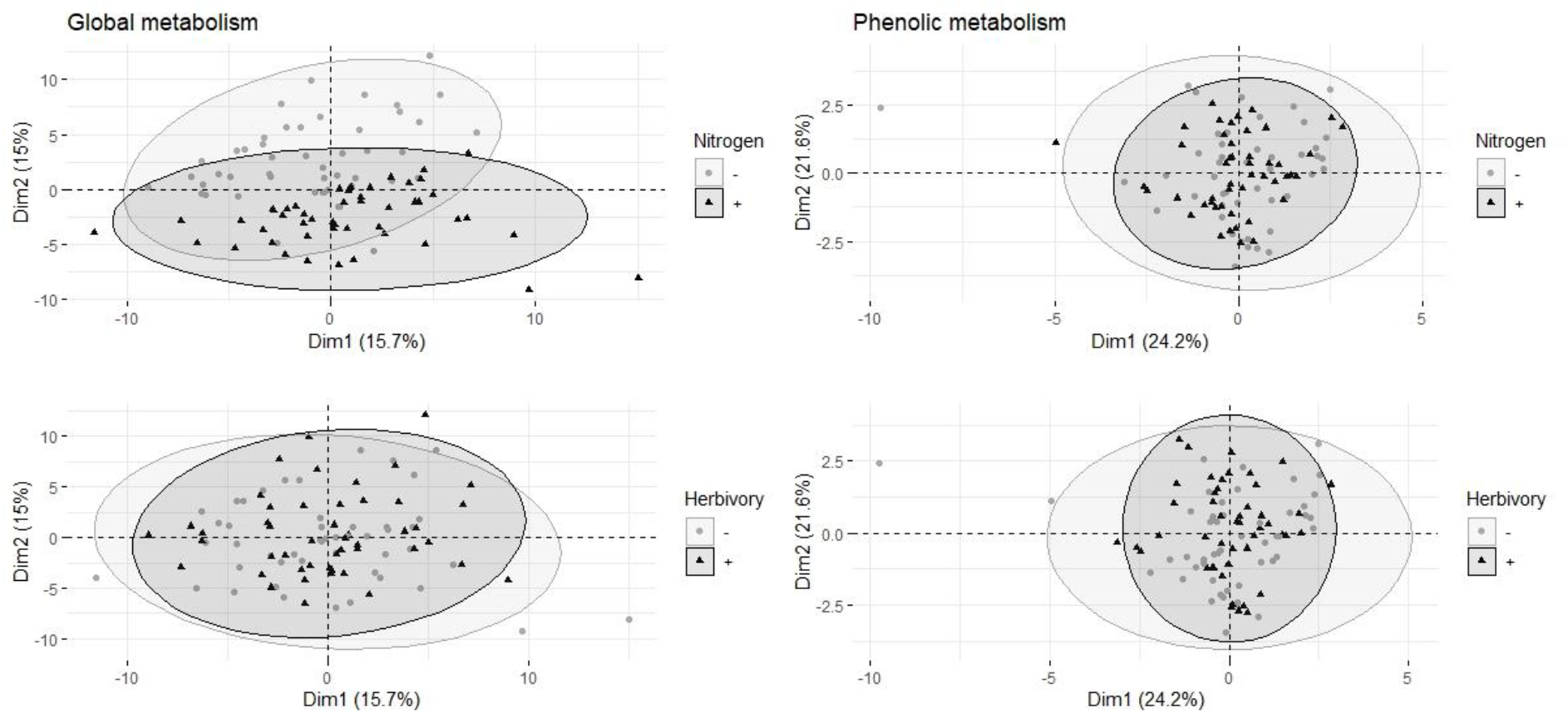
Multivariate analyses of metabolites in the leaves of aspen (*Populus tremula*) plants exposed to environmental endophytic inoculum in controlled, greenhouse-conditions: untreated control plants (C) and plants subjected to nitrogen addition (N), herbivory (H), or nitrogen addition and herbivory in combination (NH) (n = 24 plants per treatment). Shown are the Principal Component Analysis score plots of a) 114 targeted and non-targeted foliar metabolites (global analysis, data acquired using gas chromatography/mass spectrometry) and b) 11 targeted metabolites (the main salicinoid phenolic glycosides and catechin; data acquired using liquid chromatography).

The concentration of condensed tannins tended to decrease due to N and H, and for the CT precursor, catechin, the effect of N (but not H) was significant (Table 1; Fig. 2 a, b). Nitrogen treatment resulted in the lowest mean concentration and the lowest within-treatment variation in CTs was found in N treated plants (Table 1; Fig. 2 e), despite of the varied genetic background of the trees (12 clones). Overall, the effect of N on chemical characters appeared stronger compared to the effect of H (Table 1; Figs 2a-f).

**Fig. 2.**
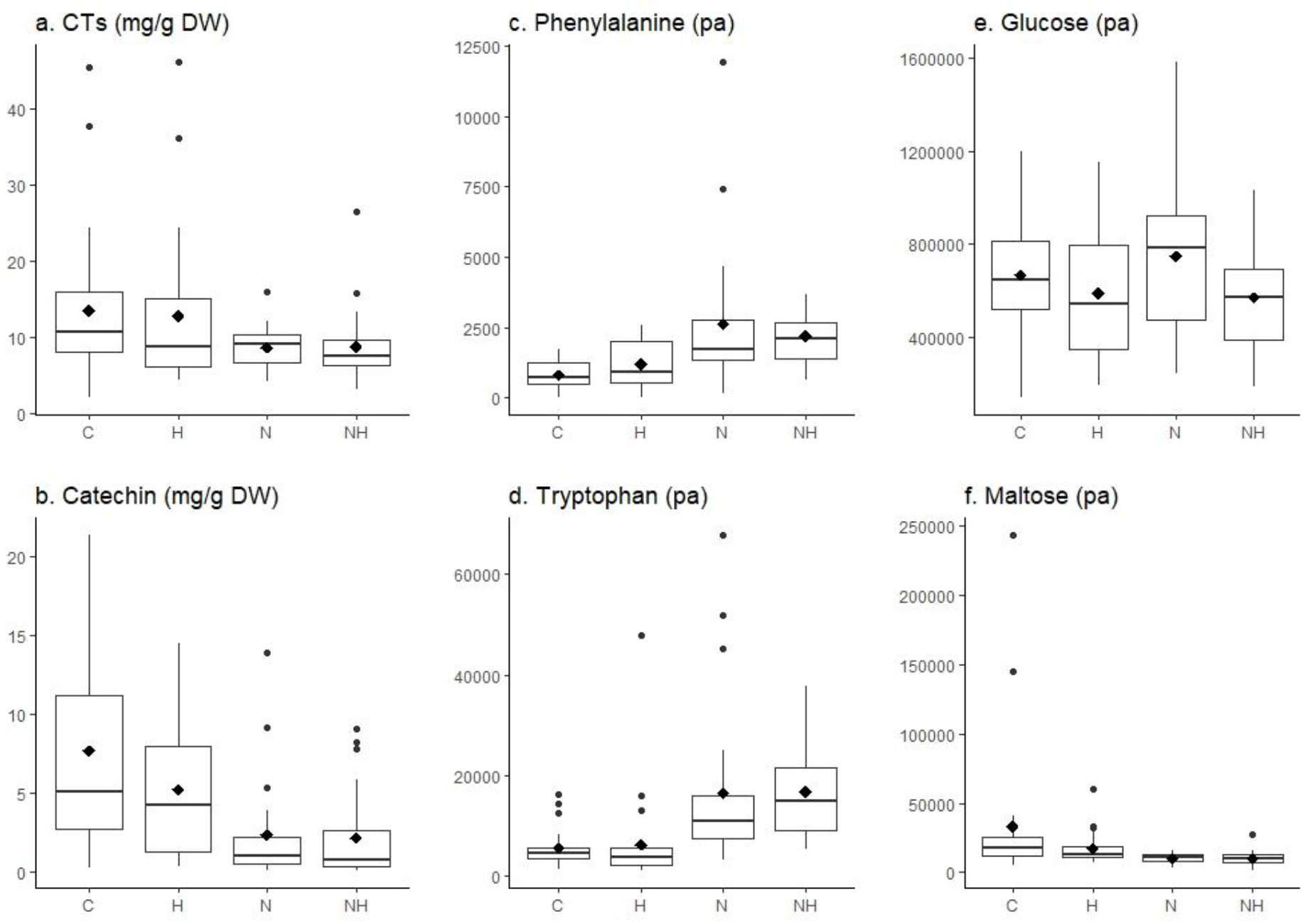
Concentration (mg g^-1^ DW) or level (measured as normalized peak area, pa) of individual chemical compounds in the leaves of aspen (*Populus tremula*) plants exposed to environmental endophytic inoculum in controlled, greenhouse-conditions: untreated control plants (C) and plants subjected to nitrogen addition (N), herbivory (H), or nitrogen addition and herbivory in combination (NH) (n = 24 plants per treatment). Shown are selected examples for phenolics (a-b: concentrations of condensed tannins and their precursor catechin), aminoacids (c-d: phenylalanine and tryptophan), and sugars (e-f: glucose, maltose). The data was acquired using liquid chromatography, except for condensed tannins which we analysed using a spectroscopy.

**Table 1.** Kruskall-Wallis statistics (chi-square, df = 3; p-value and significance α < 0.05) on treatment effects on selected phenolic compounds (condensed tannins and their precursor catechin), amino acids (phenylalanine and tryptophan) and sugars (glucose and maltose).

| **Chemicals** | **chi-square** | **p-value** | **Significance** |
| --- | --- | --- | --- |
| ***Phenolics*** |  |  |  |
| Condensed tannins | 5.73 | 0.1255 | n.s. |
| Catechin | 25.46 | 1.239e-05 | *** |
| ***Amino acids*** |  |  |  |
| Phenylalanine | 31.45 | 6.81e-07 | *** |
| Tryptophan | 42.04 | 3.928e-09 | *** |
| ***Sugars*** |  |  |  |
| Glucose | 4.86 | 0.1817 | n.s. |
| Maltose | 21.30 | 9.122e-05 | *** |

The concentrations of amino acids increased in plants receiving N treatment (see the examples in Table 1, Fig 2 a-b). Most sugars tended to be reduced by H and increased by N treatment (data for glucose is shown as an example; Table 1, Fig 2 c). A deviating pattern was found only for maltose, which was reduced by treatments, especially by N, alone or in combination with H (Table 1, Fig. 2d).

### Endophyte diversity in aspen leaves

Among the 30 MTs, 18 were identified to class (2 MTs), genus (10 MTs) or species (6 MTs) level (Table 1, Table S2). Of the identified isolates, 17 belonged to *Ascomycota* (class *Dothideomycetes, Eurotiomycetes, Hypocreales, and Sordariomycetes*) and one to *Basidiomycota* (MT25: *Rhodotorula sp*.). The three most common morphotypes across the treatments, *Ramularia* sp. (MT1), and the unidentified morphotypes MT2 and MT3 had 576 isolates (35% of all isolates), 336 (20% of all isolates) and 329 (20% of all isolates), respectively. A total of 18 MTs were captured as a maximum of three isolates (< 0.2% relative abundance) and were thus considered as rare MTs.

The sample size-based rarefaction and extrapolation curves (Fig. 3a) are predicted to accurately estimate the number of MTs per sampling unit (tree) within the double number of species when compared to the reference sample size (Chao et al. 2014). At least two fungal isolates were recovered from each plant, and the highest increase of diversity was obtained when increasing the number of sampled trees from 1 to 5 within all the treatments. The non-asymptotic rarefaction curves suggest that additional sampling efforts would yield more MTs, especially in H treatment. However, the coverage-based rarefaction and extrapolation curves (Fig. 3b) suggested that the sample size (24 trees for C and N each, and 21 for H and NH) allowed a high completeness with the method that was used: the sample coverage values were 0.95, 0.97, 0.91 and 0.93 for C, N, H, and NH, respectively.

**Fig. 3.**
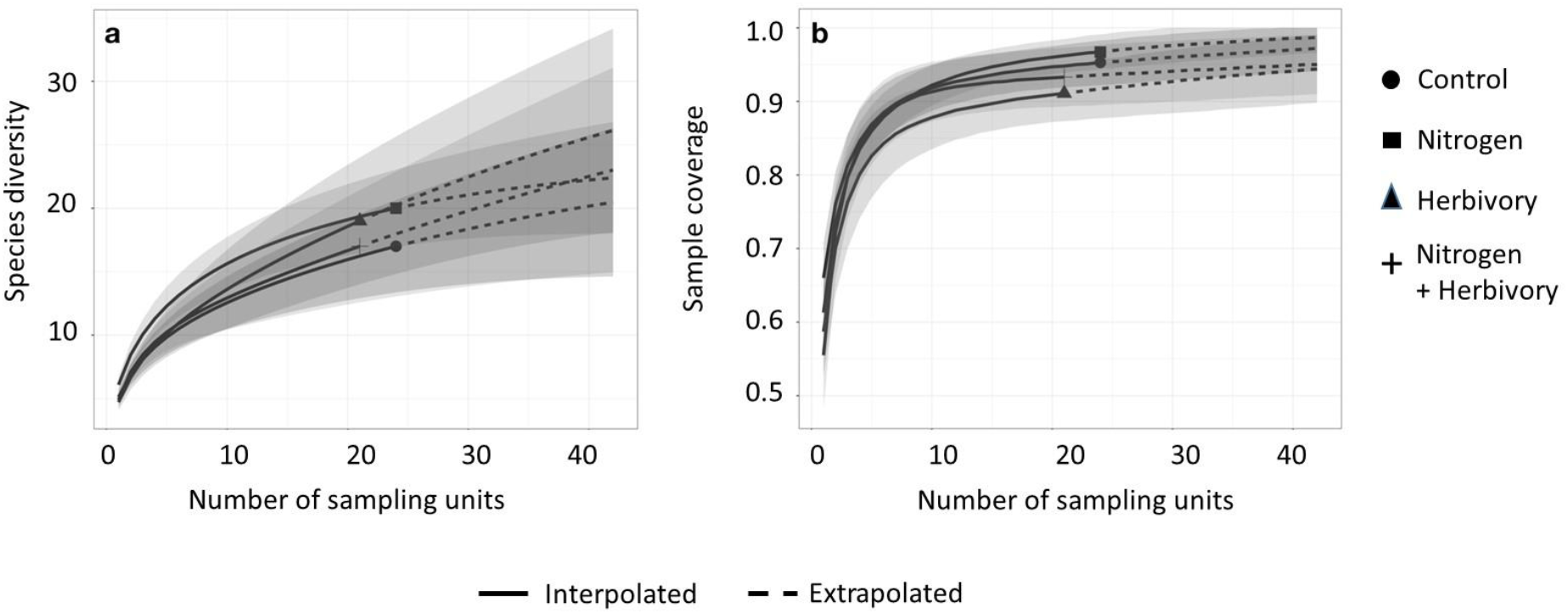
Sample-size-based (a) and (b) coverage-based rarefaction (solid line) and extrapolation (dotted line) curves comparing fungal morphotype richness in the leaves of greenhouse-cultivated aspen (*Populus tremula*) plants (n=96): untreated control plants (C) and plants subjected to nitrogen addition (N), herbivory (H), or nitrogen addition and herbivory in combination (NH) (n = 24 plants per treatment). The shaded areas represent the 95% confidence intervals. The different symbols represent the reference samples.

### Treatment effects on endophytes

The treatments caused significant differences in endophyte richness (Robust ANOVA, F=3.70, *p*=0.02) and relative abundance (Permanova, F=1.67, *p*=0.04). These differences were located between N and H treatments (*p*=0.01 for richness; and F=2.92, *p*=0.02, for abundance), while the rest of combinations did not result in significant differences. Together, the isolates belonging to the three dominating MTs made up 81% of all isolates in C plants, but their proportion tended to decrease by treatments: in N and H treatments they made up 73% of all isolates, and in NH 71%. At the same time, the relative abundance of the other (less frequent, and rare) morphotypes increased correspondingly (Table 1).

The Shannon index (H’) did not differ among the treatments (ANOVA, F=1.54, *p*=0.21), with mean (± se) and SE values 1.35 (± 0.07) for C; 1.50 (± 0.06) for N, 1.24 (± 0.10) for H; and 1.36 (± 0.08) for NH. Likewise, the Simpson index (D) values did not vary among treatments (0.68 ± 0.02 for C; 0.72 ± 0.02 for N; 0.62 ± 0.04 for H; and 0.68 ± 0.03 for NH; robust ANOVA, F=0.41, *p*=0.74), nor did the values of Pielou’s measure of evenness (J) (0.85 ± 0.02 for C; 0.84 ± 0.02 for N, 0.82 ± 0.03 for H and 0.86 ± 0.02 for NH; robust ANOVA, F=1.03, *p*=0.39).

In total, 11 MTs were shared among all four treatments (Fig 4.), of which six morphotypes were identified at least to genus level (*Ramularia* sp., *Cladosporium* sp., *Penicillium olssonii, Arthrinium* sp. and 2, *Penicillium* sp.) and five remained unknown (Table 1). Three rare MTs were recovered only from C plants, including a Dothideomycetes and a *Coleosporium* species, three others from N treated plants (a Dothideomycetes, *Physalospora scripi*, and an unknown species). Three rare MTs, including the identified *Aureobasidium* species (*A. microstictum* and *A. pullulans*) were found only in treatments that included H (Table 2).

**Fig. 4.**
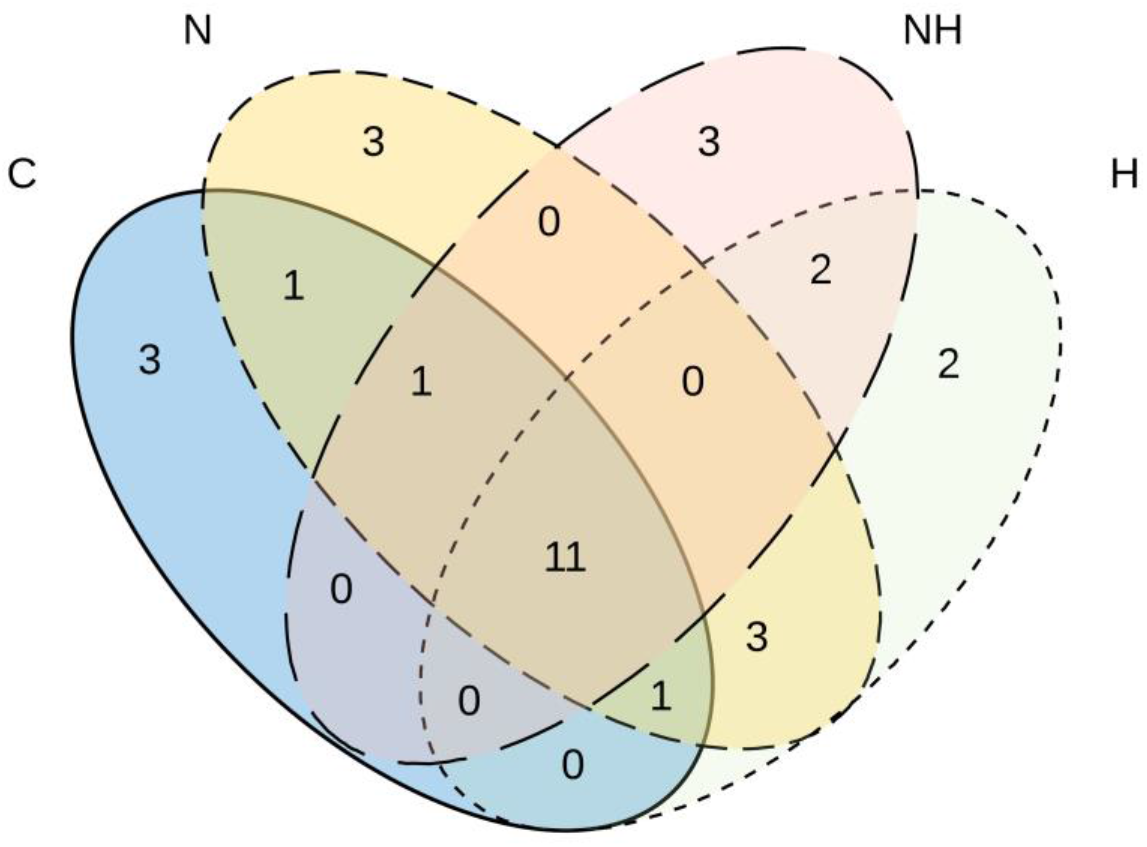
Venn diagram showing the number of unique and shared morphotypes of fungal endophytes in the leaves of greenhouse-cultivated aspen (*Populus tremula*) plants untreated control plants (C) and plants subjected to nitrogen addition (N), herbivory (H), or nitrogen addition and herbivory in combination (NH) (n = 24 plants per treatment).

**Table 2.** Morphotype identity and abundance (nr of isolates belonging to each morphotype per plant) detected in the leaves of aspen plants in different treatments treatments (control, C; herbivory, H; nitrogen fertilization, N; nitrogen fertilization and herbivory, NH).

| Morphotype | Identity | Treatment |  |  |  |
| --- | --- | --- | --- | --- | --- |
|  |  | C | N | H | NH |
| MT1 | *Ramularia sp.* | 8.83 | 6.88 | 5.14 | 4.33 |
| MT2 |  | 4.71 | 5.13 | 1.48 | 3.29 |
| MT3 | *Fusarium oxysporum* | 0.04 | 0.04 | 0.00 | 0.05 |
| MT4 |  | 0.00 | 0.04 | 0.05 | 0.00 |
| MT5 |  | 4.83 | 3.92 | 2.71 | 2.95 |
| MT6 |  | 0.17 | 0.08 | 0.05 | 0.05 |
| MT7 |  | 0.04 | 0.00 | 0.00 | 0.00 |
| MT8 | *Dothideomycetes 1* | 0.00 | 0.08 | 0.00 | 0.00 |
| MT9 | *Dothideomycetes 2* | 0.04 | 0.00 | 0.00 | 0.00 |
| MT10 |  | 0.00 | 0.00 | 0.05 | 0.05 |
| MT11 |  | 0.00 | 0.00 | 0.05 | 0.00 |
| MT12 | *Penicillium olssonii* | 0.42 | 0.67 | 0.57 | 0.81 |
| MT13 | *Cladosporium sp. 1* | 1.08 | 1.29 | 0.71 | 0.76 |
| MT14 | *Cladosporium sp. 2* | 0.04 | 0.00 | 0.00 | 0.00 |
| MT15 | *Penicillium 1* | 0.54 | 0.54 | 0.10 | 0.52 |
| MT16 |  | 0.00 | 0.04 | 0.00 | 0.00 |
| MT17 | *Penicillium brevicompactum* | 0.00 | 0.46 | 0.19 | 0.00 |
| MT18 |  | 0.88 | 0.83 | 0.38 | 0.43 |
| MT19 |  | 0.04 | 0.04 | 0.05 | 0.00 |
| MT20 | *Penicillium sp 2* | 0.00 | 0.00 | 0.00 | 0.05 |
| MT21 | *Penicillium sp. 3* | 0.04 | 0.08 | 0.00 | 0.00 |
| MT22 | *Arthrinium sp 1* | 0.54 | 0.75 | 0.52 | 0.52 |
| MT23 | *Arthrinium sp 2* | 0.08 | 0.25 | 0.05 | 0.05 |
| MT24 | *Arthrinium sp 3* | 0.00 | 0.00 | 0.05 | 0.00 |
| MT25 | *Rhodotorula sp.* | 0.00 | 0.21 | 0.14 | 0.00 |
| MT26 |  | 0.29 | 0.29 | 0.43 | 0.71 |
| MT27 | *Aureobasidium microstictum* | 0.00 | 0.00 | 0.00 | 0.14 |
| MT28 | *Aureobasidium pullulans* | 0.00 | 0.00 | 0.05 | 0.05 |
| MT29 |  | 0.00 | 0.00 | 0.00 | 0.05 |
| MT30 | *Physalospora scirpi* | 0.00 | 0.04 | 0.00 | 0.00 |
| Total nr of isolates |  | 543 | 520 | 268 | 311 |

## DISCUSSION

Nitrogen treatment induced the expected changes in the quality of aspen leaves: in particular the global metabolite analysis and condensed tannin levels highlighted the difference in the chemical profile between fertilized and non-fertilized aspen plants. A characteristic change in the global metabolite pool seemed to be due to increased concentrations of amino acids and sugars, which could reflect the expected positive effect of N on growth of the plants (Table S1, growth data). In accordance with results of several previous studies (Ibrahim et al., 2011; Bandau et al., 2015; Decker et al., 2016), we found lower concentration of CTs in leaves of fertilized plants, and N treatment also reduced the variation in CT concentration among the studied trees. Thus, nitrogen fertilization seemed to render the leaves to a more nutritious and less toxic environment (assuming antifungal effects of polyphenolics), but also a more homogenous substrate for the fungi.

In contrast to our initial expectation, however, these quantitative changes were not accompanied by a more uniform fungal community in aspen leaves when compared with the control plants. Thus, our findings did not support for the classic heterogeneity–diversity relationship hypothesis among the endophyte community in aspen leaves (i.e., the apparently more homogenous chemical environment, especially in terms of condensed tannins, was not accompanied by a lower endophyte diversity). However, recent studies suggest that heterogeneity–diversity relationships may be non-linear and more complex than expected from the niche-based perspective (Ben-Hur & Kadmon, 2020). A deeper taxonomic analysis, using culture-independent approaches, may be needed to reveal these relationships. Clearly, more detailed information about nutritional niches and functional dynamics of different endophytic fungi (Blumenstein et al. 2015; Paungfoo-Lonhienne et al., 2015) would facilitate analyses of the niche-dependent mechanisms behind the observed patterns in fungal diversity.

We expected that leaf herbivory would increase both the compositional (magnitude) and configurational (spatial) chemical heterogeneity in the leaf tissues by inducing changes in phenolic metabolism, and that this change would lead to a more complex community structure among the relatively fast growing, culturable fraction of endophyte communities. However, we did not find evidence for the expected changes in CTs or other phenolics in response to herbivory. Possibly, the impact of herbivory could have been stronger in younger leaves (a systemic induction), or after a longer feeding period. Moreover, the MT richness in H treated plants did not increase as compared to C plants. Interestingly, we found that three of the MTs were found only in connection to herbivory, supporting the view that insects may influence the fungal diversity by carrying specific fungi to the plants (Albrectsen et al., 2018). Wounding by herbivores may also open up more entry points for the ubiquitous fungi such as *Aureobasidium pullulans*, which was identified among the three H-treatment specific MTs, and is commonly found as an epiphyte and endophyte in different environments (Martín et al. 2013; Albrectsen et al., 2018).

The combined effect of N and H on the chemical traits or fungal diversity seemed to be dominated by the N effect. The longer impact time of N was likely to accentuate its effect as a stronger bottom-up force and allowed it to shape the quality of leaves as a substrate for fungi more than the short term herbivory. Fungi transmitted to plants by the feeding beetles would probably need a longer incubation time than what was possible in our study, before they can be captured using the culturing approach. It should also be noted that in our study the insects were reared in laboratory conditions and thus exposed to only a limited environmental inoculum. Therefore, the results of this study may underestimate the importance of insect-mediated facilitation of infections that occurs in natural environments.

In accordance with the universal pattern of species abundance distribution (McGill et al. 2007), we found that the culturable endophyte community in aspen leaves was composed of a few common species (the main community member *Ramularia* spp. and two unknown ones) but many rare species. The identified taxa represented *Ascomycota* commonly found as endophytes in trees, including aspen (Arnold et al., 2007; Rodriguez et al., 2009; Unterseher, 2011). The only *Basidiomycota* species identified, *Rhodotorula* sp., has been reported as a common endophyte in trees in previous studies (Unterseher, 2011; Jumpponen and Jones, 2010; Rajala et al., 2014) and it has also been linked to herbivory treatment (Albrectsen et al., 2018). In our study, however, only weak link was found between *Rhodotorula* sp. and herbivory. Intriguing was also that some common aspen endophytes, such as *Alternaria* and *Phomopsis* (Albrectsen et al., 2010; Crous and Groenewald, 2013; Martín et al., 2013), were not detected, possibly because the greenhouse acted as a filter for the environmental inocula and reduced the frequency of otherwise common endophytes. Moreover, the time point of sampling in early summer may be reflected in the community composition: the species diversity is likely to increase during the season as the infections accumulate. Among the other genera identified, *Cladosporium*, *Ramularia* (anamorph *Mycosphaerella*) and *Physalospora* have been previously described as plant pathogens (Thomma et al., 2005; Tadych et al., 2012; Videira et al., 2016), but no signs of pathogen attacks were found on the leaves, suggesting that the conditions in our experiment supported endophytic lifestyle of these and other fungi.

In conclusion, our results indicate that the culturable fraction of fungal endophyte community in aspen leaves is rather stable against the directional alterations induced in the chemical “landscape” within the leaves by nitrogen fertilization or disturbances due to herbivory. This is in line with the results of the study by Christian et al. (2016) who reported high resistance among endophyte communities against biotic and anthropogenic perturbations. However, more research is needed to better comprehend the nutritional requirements of endophytes, and the signals influencing the dynamics of the microbial communities in the endosphere. It should also be acknowledged that different fertilization levels and sampling times could result in stronger responses in the chemical environment and fungal communities. Thus, our results should be generalized with caution. While our results did not agree with the expectations we made based on the heterogeneity-diversity relationship hypothesis, we propose that by implementing theoretical frameworks of community ecology it is possible to gain new insights into the processes and traits driving the structure and functions of endophyte communities in perennial plants, such as forest trees.

## Acknowledgments

We thank Sylvia S. Chen for assistance in the experiments. The research was supported by FORMAS, a Swedish Research Council for Sustainable Development (2012-01358 and 2016-00907) to JW, and The Royal Swedish Academy of Agriculture and Forestry to MA.

## References

Abreu IN, Ahnlund M, Moritz T, Albrectsen BR (2011). UHPLC-ESI/TOFMS determination of salicylate-like phenolic gycosides in *Populus tremula* leaves. J. Chem. Ecol. 37:857–870. doi:10.1007/s10886-011-9991-7.

Agostinelli M, Cleary M, Martín JA, Albrectsen BR, Witzell J (2018). Pedunculate oaks *(Quercus robur* L.) differing in vitality as reservoirs for fungal biodiversity. Front. Microbiol. 9:1758. doi:10.3389/fmicb.2018.01758.

Albrectsen BR, Witzell J (2012). “Disentangling functions of fungal endophytes in forest trees,” in Fungi: types, environmental impact and role in disease, eds. A. Paz Silva and M. Sol (Huntington, NY: Nova Science Publishers), pp. 235–246.

Albrectsen BR, Björkén L, Varad A, Hagner Å, Wedin M, Karlsson J, Jansson S (2010). Endophytic fungi in European aspen (*Populus tremula*) leaves - diversity, detection, and a suggested correlation with herbivory resistance. Fungal Div. 41:17–28. doi:10.1007/s13225-009-0011-y.

Albrectsen BR, Siddique AB, Decker VHG, Unterseher M, Robinson KM (2018). Both plant genotype and herbivory shape aspen endophyte communities. Oecologia 187:535–545. doi:10.1007/s00442-018-4097-3.

Arnold AE, Henk DA, Eells RL, Lutzoni F, Vilgalys R (2007). Diversity and phylogenetic affinities of foliar fungal endophytes in loblolly pine inferred by culturing and environmental PCR. Mycologia 99: 185–206. doi:10.3852/mycologia.99.2.185.

Badotti F, de Oliveira FS, Garcia CF, (2017) Effectiveness of ITS and sub-regions as DNA barcode markers for the identification of Basidiomycota (Fungi). BMC Microbiol. 17, 42 doi: 10.1186/s12866-017-0958-x

Bailey JK, Deckert R, Schweitzer JA, Rehill BJ, Lindroth RL, Gehring C, Whitham TG (2005). Host plant genetics affect hidden ecological players: links among *Populus,* condensed tannins, and fungal endophyte infection. Can. J. Bot. 83:356–361. doi:10.1139/b05-008.

Bandau F, Decker VHG, Gundale MJ, Albrectsen BR (2015). Genotypic tannin levels in *Populus tremula* impact the way nitrogen enrichment affects growth and allocation responses for some traits and not for others. PLoS ONE 10(10): e0140971. doi:10.1371/journal.pone.0140971.

Ben-Hur E, Kadmon R (2020). Heterogeneity-diversity relationships in sessile organisms: a unified framework. Ecol. Lett. 23:193–207. doi:10.1111/ele.13418.

Blacutt A, Ginnan N, Dang T, Bodaghi S, Vidalakis G, Ruegger P, Peacock B, Viravathana P, Vieira FC, Drozd C, Jablonska B, Borneman J, McCollum G, Cordoza J, Meloch J, Berry V, Salazar LL, Maloney KN, Rolshausen PE, Roper MC (2020). An in vitro pipeline for screening and selection of citrus-associated microbiota with potential anti-”*Candidatus Liberibacter asiaticus”* properties. Appl Environ Microbiol 86:e02883–19. https://doi.org/10.1128/AEM.02883-19.

Blumenstein K, Albrectsen BR, Martín JA, Hultberg M, Sieber TN, Helander M, Witzell J (2015). Nutritional niche overlap potentiates the use of endophytes in biocontrol of a tree disease. BioControl 60: 655–667. doi:10.1007/s10526-015-9668-1.

Busby PE, Peay KG, Newcombe G (2016). Common foliar fungi of *Populus trichocarpa* modify *Melampsora* rust disease severity. New Phytol. 209:1681–1692. doi:10.1111/nph.13742.

Carroll G (1995). Forest endophytes: pattern and process. Can. J. Bot. 73:1316–1324. doi:10.1139/b95-393.

Chao A, Gotelli NJ, Hsieh TC, Sander EL, Ma KH, Colwell RK, Ellison AM (2014). Rarefaction and extrapolation with Hill numbers: a framework for sampling and estimation in species diversity studies. Ecol. Monogr. 84: 45–67.

Christian N, Sullivan C, Visser ND, Clay K (2016). Plant host and geographic location drive endophyte community composition in the face of perturbation. Microb Ecol. 72:621–32. doi:10.1007/s00248-016-0804-y.

Clay, K (1988). Fungal endophytes of grasses: a defensive mutualism between plants and fungi. Ecology 69:10–16.

Coblentz KE, Van Bael SA (2013). Field colonies of leaf-cutting ants select plant materials containing low abundances of endophytic fungi. Ecosphere 4:1–10. doi:10.1890/ES13-00012.1.

Colwell RK, Chao A, Gotelli NJ, Lin S-Y, Mao CX, Chazdon RL, Longino JT (2012). Models and estimators linking individual-based and sample-based rarefaction, extrapolation and comparison of assemblages. J. Plant Ecology 5: 3–21.

Constabel CP, Yip L, Patton JJ, Christopher ME (2000). Polyphenol oxidase from hybrid poplar. Cloning and expression in response to wounding and herbivory. Plant Physiol. 124:285–295. doi:10.1104/pp.124.1.285

Crous PW, Groenewald JZ (2013). A phylogenetic re-evaluation of *Arthrinium*. IMA Fungus 4: 133–54. doi:10.5598/imafungus.2013.04.01.13.

Decker VHG, Bandau F, Gundale MJ, Cole CT, Albrectsen BR (2016). Aspen phenylpropanoid genes’ expression levels correlate with genets’ tannin richness and vary both in responses to soil nitrogen and associations with phenolic profiles. Tree Physiol. 37: 270–279. doi:10.1093/treephys/tpw118.

Edenius L, Mikusinski G, Witzell J, Bergh J (2012) Effects of repeated fertilization of young Norway spruce on phenolics and arthropods: implications for insectivorous birds’ food resources. For. Ecol. Manage. 277: 38–45. doi:10.1016/j.foreco.2012.04.021.

Faeth SH., Sullivan TJ (2003). Mutualistic asexual endophytes in a native grass are usually parasitic. Am. Nat. 161: 310–325. doi:10.1086/345937.

Gomes T, Pereira JA, Benhadi J, Lino-Neto T, Baptista P (2018). Endophytic and epiphytic phyllosphere fungal communities are shaped by different environmental factors in a Mediterranean ecosystem. Microb Ecol 76:668–679. doi:10.1007/s00248-018-1161-9.

Gouda S, Das G, Sen SK, Shin H-S, Patra JK (2016). Endophytes: a treasure house of bioactive compounds of medicinal importance. Front. Microbiol. 7: 1–8. doi:10.3389/fmicb.2016.01538.

Gourlay G, Constabel P (2019). Condensed tannins are inducible antioxidants and protect hybrid poplar against oxidative stress. Tree Physiol. 39: 345–355, doi:10.1093/treephys/tpy143

Hardoim PR, van Overbeek LS, Berg G, Pirttilä AM, Compant S, Campisano A, Döring M, Sessitsch A (2015). The hidden world within plants: ecological and evolutionary considerations for defining functioning of microbial endophytes. Microbiol. Mol. Biol. Rev. 79: 293–320. doi:10.1128/MMBR.00050-14.

Hsieh TC, Ma KH, Chao A (2016). iNEXT: An R package for rarefaction and extrapolation of species diversity (Hill numbers). Meth. Ecol. Evol. 7: 1451–1456. doi:10.1111/2041-210X.12613.

Ibrahim MH, Jaafar HZE, Rahmat A, Rahman ZA (2011). Effects of nitrogen fertilization on synthesis of primary and secondary metabolites in three varieties of Kacip Fatimah *(Labisia pumila* Blume). Int. J. Mol. Sci. 12: 5238–54. doi:10.3390/ijms12085238.

Jonsson P, Johansson AI, Gullberg J, Trygg J, A J, Grung B., Marklund S, Sjöström M, Antti H, Moritz T (2005). High-throughput data analysis for detecting and identifying differences between samples in GC/MS-based metabolomic analyses. Anal. Chem. 77: 5635–5642. doi:10.1021/ac050601e.

Jumpponen, A, Jones KL (2009). Massively parallel 454 sequencing indicates hyperdiverse fungal communities in temperate *Quercus macrocarpa* phyllosphere. New Phytol. 184:438–448. doi:10.1111/j.1469-8137.2009.02990.x.

Jumpponen A, Jones KL (2010). Seasonally dynamic fungal communities in the *Quercus macrocarpa* phyllosphere differ between urban and nonurban environments. New Phytol. 186:496–513. doi:10.1111/j.1469-8137.2010.03197.x.

Keefover-Ring K, Ahnlund M, Abreu IN, Jansson S, Moritz T, Albrectsen BR (2014). No evidence of geographical structure of salicinoid chemotypes within *Populus tremula*. PLoS ONE 9(10): e107189. doi:10.1371/journal.pone.0107189.

Lê S, Josse J, Husson F (2008). FactoMineR: An R Package for Multivariate Analysis. J. Stat. Soft. 25:1–18. doi. 10.18637/jss.v025.i01.

Lewinsohn D, Lewinsohn E, Bertagnolli CL, Patridge AD (1994). Blue-stain fungi and their transport structures on the Douglas-fir beetle. Can. J. For. Res. 24: 2275–2283. doi:10.1139/x94-292.

Martín JA, Witzell J, Blumenstein K, Rozpedowska E, Helander M, Sieber TN, Gil L (2013). Resistance to Dutch elm disease reduces presence of xylem endophytic fungi in elms *(Ulmus* spp.). PLoS ONE 8(2): e56987. doi:10.1371/journal.pone.0056987.

Martinez Arbizu P (2019). pairwiseAdonis: Pairwise multilevel comparison using adonis. R package version 0.3.

Massad TJ, Trumbore SE, Ganbat G, Reichelt M, Unsicker S, Boeckler A, Gleixner G, Gershenzon J, Ruehlow S (2014) An optimal defense strategy for phenolic glycoside production in *Populus trichocarpa* – isotope labeling demonstrates secondary metabolite production in growing leaves. New Phytol. 203:607–619. doi:10.1111/nph.12811.

McGill BJ, Etienne RS, Gray JS, Alonso D, Anderson MJ, Benecha HK, Dornelas M, Enquist BJ, Green JL, He F, Hurlbert AH, Magurran AE, Marquet PA, Maurer BA, Ostling A, Soykan CU, Ugland KI, White EP (2007). Species abundance distributions: moving beyond single prediction theories to integration within an ecological framework. Ecol. Lett. 10:995–1015. doi:10.1111/j.1461-0248.2007.01094.x.

Nisa H, Kamili AN, Nawchoo IA, Shafi S, Shameem N, Bandh SA (2015). Fungal endophytes as prolific source of phytochemicals and other bioactive natural products: A review. Microb. Pathog. 82:50–59. doi:10.1016/j.micpath.2015.04.001.

Nohrstedt H-Ö (2001). Response of coniferous forest ecosystems on mineral soils to nutrient additions: a review of Swedish experiences. Scand. J. For. Res. 16:555–573. doi:10.1080/02827580152699385.

Oksanen J, Blanchet FG, Kindt R, Legendre P, Minchin P, O’Hara R, et al. (2016). vegan: Community Ecology Package. R package version 2.4-0.

Paungfoo-Lonhienne C, Yeoh YK, Kasinadhuni NRP, Lonhienne TGA, Robinson N, Hugenholtz P, Ragan MA, Schmidt S (2015). Nitrogen fertilizer dose alters fungal communities in sugarcane soil and rhizosphere. Sci. Rep. 5, 8678. doi:10.1038/srep08678.

Petrini O, Sieber TN, Toti L, Viret O (1993). Ecology, metabolite production, and substrate utilization in endophytic fungi. Nat. Toxins 1: 185–196. doi:10.1002/nt.2620010306.

Porter JK, Bacon CW, Cutler HG, Arrendale RF, Robbins JD (1985). In vitro auxin production by *Balansia epichloë*. Phytochemistry 24:1429–1431. doi:10.1016/S0031-9422(00)81037-7.

R Development Core Team (2016). R: A language and environment for statistical computing. Vienna, Austria: R Foundation for Statistical Computing Available at: https://www.r-project.org/.

Ragazzi A, Moricca S, Capretti P, Dellavalle I, Turco E (2003). Differences in composition of endophytic mycobiota in twigs and leaves of healthy and declining *Quercus* species in Italy. For. Pathol. 33: 31–38. doi:10.1046/j.1439-0329.2003.3062003.x.

Rajala T, Velmala SM, Vesala R, Smolander A, Pennanen T (2014). The community of needle endophytes reflects the current physiological state of Norway spruce. Fungal Biol. 118: 309–15. doi:10.1016/j.funbio.2014.01.002.

Rice AV, Thormann MN, Langor DW (2007). Mountain pine beetle associated blue-stain fungi cause lesions on jack pine, lodgepole pine, and lodgepole × jack pine hybrids in Alberta. Can. J. Bot. 85: 307–315. doi:10.1139/B07-014.

Robinson KM, Ingvarsson PK, Jansson S, Albrectsen BR (2012). Genetic variation in functional traits influences arthropod community composition in Aspen (*Populus tremula* L.). PLOS ONE 7(5): e37679, doi:0.1371/journal.pone.0037679.

Rodriguez RJ, White JF, Arnold AE, Redman RS (2009). Fungal endophytes: diversity and functional roles. New Phytol. 182: 314–30. doi:10.1111/j.1469-8137.2009.02773.x.

Salminen JP, Karonen M (2011). Chemical ecology of tannins and other phenolics: we need a change in approach. Funct. Ecol. 25: 325–338. doi:10.1111/j.1365-2435.2010.01826.x

Sampedro L, Moreira X, Zas R (2011). Costs of constitutive and herbivore-induced chemical defences in pine trees emerge only under low nutrient availability. J. Ecol. 99:818–827. doi:10.1111/j.1365-2745.2011.01814.x.

Schauer N, Steinhauser D, Strelkov S, Schomburg D, Allison G, Moritz T, Lundgren K, Roessner-Tunali U, Forbes MG, Willmitzer L, Fernie AR, Kopka J (2005). GC–MS libraries for the rapid identification of metabolites in complex biological samples. FEBS Letters, 579. doi:10.1016/j.febslet.2005.01.029.

Shade A, Handelsman J (2012). Beyond the Venn diagram: the hunt for a core microbiome. Environ. Microbiol. 14, 4–12. doi:10.1111/j.1462-2920.2011.02585.x.

Sieber TN (2007). Endophytic fungi in forest trees: are they mutualists? Fungal Biol. Rev. 21, 75–89. doi:10.1016/j.fbr.2007.05.004.

Tadych M, Bergen MS, Johnson-Cicalese J, Polashock JJ, Vorsa N, White JF (2012). Endophytic and pathogenic fungi of developing cranberry ovaries from flower to mature fruit: diversity and succession. Fungal Div. 54: 101–116. doi:10.1007/s13225-012-0160-2.

Terhonen E, Blumenstein K, Kovalchuk A, Asiegbu F (2019). Forest tree microbiomes and associated fungal endophytes: Functional roles and impact on forest health. Forests 10: 42. doi:10.3390/f10010042

Thomma BPHJ, Van Esse HP, Crous PW, De Wit PJGM (2005). *Cladosporium fulvum* (syn. *Passalora fulva*), a highly specialized plant pathogen as a model for functional studies on plant pathogenic Mycosphaerellaceae. Mol. Plant Pathol. 6: 379–393. doi:10.1111/j.1364-3703.2005.00292.x.

Ullah C, Unsicker SB, Fellenberg C, Constabel P, Schmidt A, Gershenzon J, Hammerbacher A (2017). Flavan-3-ols are an effective chemical defense against rust infection. Plant Physiol.;175:1560–1578. doi:10.1104/pp.17.00842.

Unterseher M (2011). Diversity of fungal endophytes in temperate forest trees. In: Endophytes of Forest Trees: Biology and Applications, eds. AM Pirttilä and AC Frank (Springer, Dordrecht), 31–46. doi:10.1007/978-94-007-1599-8_2.

Unterseher M, Siddique AB, Brachmann A, Peršoh D (2016). Diversity and composition of the leaf mycobiome of beech *(Fagus sylvatica)* are affected by local habitat conditions and leaf biochemistry. PLoS ONE 11(4): e0152878. doi/10.1371/journal.pone.0152878.

Vellend M (2010). Conceptual synthesis in community ecology. Quart. Rev. Biol. 85:183–206

Videira SIR, Groenewald JZ, Braun U, Shin HD, Crous PW (2016). All that glitters is not *Ramularia*. Stud. Mycol. 83:49–163. doi:10.1016/J.SIMYCO.2016.06.001.

White TJ, Bruns T, Lee S, Taylor J (1990). Amplification and direct sequencing of fungal ribosomial RNA genes for phyologenetics. In: PCR Protocols: a guide to methods and applicaitions, ed. TJ Innis, MA, Gelfand DH, Sninsky JJ. Academic Press Inc., New York, pp 315–322.

Whittaker RH (1972). Evolution and measurement of species diversity. Taxon 21: 213–251.

Wilson M, Lindow SE (1994). Coexistence among epiphytic bacterial populations mediated through nutritional resource partitioning. Appl. Environ. Microbiol. 60: 4468–77.

Witzell J, Shevtsova A (2004). Nitrogen-induced changes in phenolics of *Vaccinium myrtillus* - implications for interaction with a parasitic fungus. J. Chem. Ecol. 10:1919–1938. doi:10.1023/b:joec.0000045587.75128.a4.

Witzell J, Martín JA (2008). Phenolic metabolites in the resistance of northern forest trees to pathogens — past experiences and future prospects. Can. J. For. Res. 38:2711–2727. doi:10.1139/X08-112.

Witzell J, Martín JA (2018). Endophytes and forest health. In: Endophytes of forest trees. Eds. AM Pirttilä and CA Frank. Forestry Sciences 86. Springer International Publishing AG, Springer Nature, pp 261–282. doi:10.1007/978-3-319-89833-9_12.

Witzell J, Martín JA, Blumenstein K (2014) Ecological aspects of endophyte-based biocontrol of forest diseases. In: VC Verma & AC Gange (eds.), Advances in Endophytic Research, pp. 321–333. Springer India. doi:10.1007/978-81-322-1575-2_17.

