## Supplemental Tables 1 to 3 for "Aspen leaves as a “chemical landscape” for fungal endophyte diversity – Can nitrogen and herbivory shape the community composition in controlled conditions?"

**Supplementary Tables**

**Table S1**. Growth responses to treatments (C=control, N = nitrogen fertilization; H=herbivory, NH= combined N and H treatment; values=mean ± s.e). Different letters indicate statistic differences according to the Tukey HSD posthoc test, CI=95%.

| **Response** | **Treatment** | | | | |
|---|---|---|---|---|---|
| Above (g) | C | 19.12 | ± | 1.03 | b |
| | N | 32.54 | ± | 1.36 | a |
| | H | 17.28 | ± | 1.04 | b |
| | NH | 32.54 | ± | 1.46 | a |
| Height (cm) | C | 149.19 | ± | 5.25 | b |
| | N | 176.98 | ± | 4.14 | a |
| | H | 137.24 | ± | 4.21 | b |
| | NH | 166.63 | ± | 3.86 | a |
| Leaf area ($mm^2$) | C | 274.81 | ± | 24.41 | b |
| | N | 391.82 | ± | 25.09 | a |
| | H | 329.78 | ± | 53.13 | ab |
| | NH | 353.33 | ± | 31.97 | ab |
| Beetle weight (mg) | H | 68.54 | ± | 1.21 | a |
| | NH | 72.77 | ± | 2.58 | a |

**Table S2** Morphotypes and putative taxon correspondence with the max score, query coverage, max identity, closest GenBank match.

| MT | Putative taxon | Max score | Query coverage % | Max identity % | Closest GenBank Match |
|---|---|---|---|---|---|
| **MT1** | *Ramularia* sp. | 961 | 100 | 99 | KJ504798 |
| **MT3** | *Fusarium oxysporum* | 987 | 100 | 100 | KY587310 |
| **MT8** | *Dothideomycetes1* | 913 | 100 | 97 | JX010732 |
| **MT9** | *Dothideomycetes2* | 743 | 98 | 96 | FJ515608 |
| **MT12** | *Penicillium olsonii* | 1038 | 96 | 100 | DQ117963 |
| **MT13** | *Cladosporium* sp. *1* | 1014 | 99 | 100 | KP701958 |
| **MT14** | *Cladosproium* sp. *2* | 1016 | 99 | 100 | KU182498 |
| **MT15** | *Penicillium* sp. *1* | 1051 | 100 | 99 | AY373899 |
| **MT17** | *Penicillium brevicompactum* | 1068 | 99 | 100 | JN246028 |
| **MT20** | *Penicillium* sp. *2* | 1064 | 100 | 99 | KM520354 |
| **MT21** | *Penicillium* sp. *3* | 998 | 100 | 100 | HG530747 |
| **MT22** | *Arthrinium* sp. *1* | 1086 | 94 | 100 | KT715716 |
| **MT23** | *Arthrinium* sp. *2* | 1053 | 99 | 99 | GU566268 |
| **MT24** | *Arthrinium* sp. *3* | 1133 | 100 | 99 | KF144901 |
| **MT25** | *Rhodotorula* sp. | 1103 | 100 | 99 | AF444610 |
| **MT27** | *Aureobasidium microstictum* | 1072 | 100 | 99 | EU167608 |
| **MT28** | *Aureobasidium pullulans* | 1007 | 100 | 99 | JQ235063 |
| **MT30** | *Physalospora scirpi* | 867 | 86 | 99 | KP698184 |

**Table S3.** Diversity index values (H= Shannon, D=Simpson, J=Pielou's evenness) in replicate plants per treatments (C=control, N = nitrogen fertilization; H=herbivory, NH= combined N and H treatment).

| Treatment | Genotype | H | D | J |
|---|---|---|---|---|
| C | GT_100 | 1.348 | 0.676 | 0.753 |
| C | GT_100 | 1.490 | 0.758 | 0.926 |
| C | GT_26 | 0.691 | 0.498 | 0.997 |
| C | GT_26 | 1.297 | 0.651 | 0.806 |
| C | GT_4 | 1.602 | 0.766 | 0.894 |
| C | GT_4 | 0.935 | 0.579 | 0.851 |
| C | GT_41 | 1.488 | 0.747 | 0.925 |
| C | GT_41 | 1.223 | 0.680 | 0.882 |
| C | GT_51 | 1.840 | 0.809 | 0.885 |
| C | GT_51 | 1.505 | 0.760 | 0.935 |
| C | GT_6 | 2.008 | 0.857 | 0.965 |
| C | GT_6 | 1.335 | 0.635 | 0.686 |
| C | GT_69 | 2.051 | 0.868 | 0.986 |
| C | GT_69 | 1.161 | 0.656 | 0.837 |
| C | GT_7 | 1.037 | 0.630 | 0.944 |
| C | GT_7 | 1.004 | 0.484 | 0.624 |
| C | GT_72 | 1.566 | 0.784 | 0.973 |
| C | GT_72 | 1.356 | 0.712 | 0.842 |
| C | GT_79 | 1.270 | 0.693 | 0.916 |
| C | GT_79 | 1.133 | 0.584 | 0.704 |
| C | GT_92 | 1.402 | 0.719 | 0.871 |
| C | GT_92 | 0.874 | 0.451 | 0.631 |
| C | GT_98 | 1.590 | 0.770 | 0.888 |
| C | GT_98 | 1.181 | 0.595 | 0.734 |
| N | GT_100 | 1.163 | 0.620 | 0.723 |
| N | GT_100 | 1.871 | 0.821 | 0.900 |
| N | GT_26 | 1.327 | 0.646 | 0.740 |
| N | GT_26 | 1.642 | 0.777 | 0.916 |
| N | GT_4 | 2.095 | 0.861 | 0.953 |
| N | GT_4 | 1.500 | 0.740 | 0.837 |
| N | GT_41 | 0.955 | 0.508 | 0.689 |
| N | GT_41 | 1.479 | 0.756 | 0.919 |
| N | GT_51 | 0.944 | 0.570 | 0.859 |
| N | GT_51 | 1.413 | 0.670 | 0.789 |
| N | GT_6 | 1.735 | 0.793 | 0.892 |
| N | GT_6 | 1.569 | 0.755 | 0.876 |
| N | GT_69 | 2.014 | 0.859 | 0.969 |
| N | GT_69 | 1.851 | 0.814 | 0.890 |
| N | GT_7 | 1.103 | 0.616 | 0.796 |
| N | GT_7 | 1.318 | 0.656 | 0.736 |

| | | | | |
|---|---|---|---|---|
| N | GT_72 | 1.361 | 0.626 | 0.699 |
| N | GT_72 | 1.457 | 0.742 | 0.905 |
| N | GT_79 | 1.307 | 0.676 | 0.812 |
| N | GT_79 | 1.703 | 0.747 | 0.819 |
| N | GT_92 | 1.721 | 0.774 | 0.827 |
| N | GT_92 | 1.165 | 0.609 | 0.724 |
| N | GT_98 | 1.430 | 0.715 | 0.798 |
| N | GT_98 | 1.488 | 0.745 | 0.924 |
| H | GT_100 | 1.802 | 0.798 | 0.866 |
| H | GT_100 | 1.721 | 0.781 | 0.885 |
| H | GT_26 | 1.471 | 0.740 | 0.914 |
| H | GT_26 | 0.349 | 0.198 | 0.503 |
| H | GT_4 | 0.410 | 0.245 | 0.592 |
| H | GT_4 | 0.689 | 0.328 | 0.497 |
| H | GT_41 | 0.950 | 0.560 | 0.865 |
| H | GT_51 | 1.514 | 0.768 | 0.940 |
| H | GT_51 | 1.040 | 0.625 | 0.946 |
| H | GT_6 | 2.062 | 0.858 | 0.938 |
| H | GT_69 | 1.182 | 0.654 | 0.853 |
| H | GT_69 | 1.494 | 0.750 | 0.928 |
| H | GT_7 | 1.303 | 0.665 | 0.810 |
| H | GT_72 | 1.154 | 0.612 | 0.832 |
| H | GT_72 | 0.684 | 0.370 | 0.622 |
| H | GT_79 | 1.295 | 0.645 | 0.804 |
| H | GT_79 | 0.898 | 0.540 | 0.817 |
| H | GT_92 | 1.400 | 0.714 | 0.870 |
| H | GT_92 | 1.330 | 0.722 | 0.959 |
| H | GT_98 | 1.610 | 0.727 | 0.827 |
| H | GT_98 | 1.720 | 0.810 | 0.960 |
| NH | GT_100 | 1.319 | 0.678 | 0.819 |
| NH | GT_100 | 1.534 | 0.737 | 0.856 |
| NH | GT_26 | 0.637 | 0.444 | 0.918 |
| NH | GT_4 | 0.956 | 0.571 | 0.870 |
| NH | GT_4 | 1.330 | 0.722 | 0.959 |
| NH | GT_51 | 1.988 | 0.838 | 0.905 |
| NH | GT_51 | 1.748 | 0.816 | 0.976 |
| NH | GT_6 | 1.461 | 0.747 | 0.908 |
| NH | GT_6 | 1.677 | 0.790 | 0.936 |
| NH | GT_69 | 0.857 | 0.436 | 0.618 |
| NH | GT_69 | 1.298 | 0.704 | 0.937 |
| NH | GT_7 | 0.868 | 0.500 | 0.790 |
| NH | GT_7 | 1.667 | 0.781 | 0.931 |
| NH | GT_72 | 0.828 | 0.512 | 0.753 |
| NH | GT_72 | 1.378 | 0.698 | 0.856 |

| | | | | |
|---|---|---|---|---|
| NH | GT_79 | 1.210 | 0.630 | 0.752 |
| NH | GT_79 | 1.383 | 0.626 | 0.711 |
| NH | GT_92 | 1.710 | 0.806 | 0.955 |
| NH | GT_92 | 1.681 | 0.768 | 0.864 |
| NH | GT_98 | 1.465 | 0.741 | 0.910 |
| NH | GT_98 | 1.666 | 0.744 | 0.856 |
